# Evaluating the effects of arousal and emotional valence on performance of racing greyhounds

**DOI:** 10.1101/831552

**Authors:** Melissa Starling, Anthony Spurrett, Paul McGreevy

## Abstract

The racing greyhound industry in Australia has come under scrutiny in recent years due to animal welfare concerns, including so-called behavioural wastage whereby physically sound greyhounds are removed from the racing industry because of poor performance. The non-medical reasons why greyhounds perform poorly at the racetrack are not well understood, but may include insufficient reinforcement for racing, or negative affective states associated with the context of racing. This study sought evidence for the affective states of greyhounds (n=525) at race meets and associations of those states with performance. It collected demographic, behavioural and performance data, along with infrared thermographic images of greyhounds at race-meets to investigate whether arousal influenced performance. It also collected behavioural data in the catching pen at the completion of races to examine possible evidence of frustration that may reflect sub-optimal behavioural reinforcement.

Linear regression models were built to determine factors affecting greyhound performance. Increasing mean eye temperature after the race and increasing greyhound age both had a statistically significant, negative effect on performance. The start box number also had a significant effect, with boxes 4, 5 and 7 having a negative effect on performance. There was a significant effect of track on mean eye temperatures before and after the race, suggesting that some tracks may be inherently more stressful for greyhounds than others. Behaviours that may indicate frustration in the catching pen were extremely common at two tracks, but much less common at the third, where play objects in motion were used to draw greyhounds into the catching pen. The study provides evidence for the use of eye temperature in predicting performance, guidance for assessment of poor performance in greyhounds and suggested approaches to the management of frustration in racing greyhounds.

## Introduction

Greyhound racing in Australia is a sport supported largely by revenue from betting on the outcome of races. As such, there may be pressure on owners of racing greyhounds to race their dogs as often as they are physically able so that race winnings support the greyhounds’ upkeep and further racing activities. Recent scrutiny into the greyhound racing industry in Australia, and in particular, the state of New South Wales (NSW), has raised questions about the level of so-called wastage of greyhounds within the racing industry, which is where greyhounds are discarded from racing because they do not perform the task they were bred for, i.e., racing [1]. Wastage can be physical e.g., from lameness, or behavioural e.g., from relative disinterest in running. The ultimate fate of discarded dogs is unknown, but may include rehoming to a pet home, being retired but retained by the original owner or trainer, or euthanasia. Failing to chase a lure is considered a form of behavioural wastage, which is where otherwise physically healthy and sound animals are removed from a role because they are unable to perform it adequately due to behavioural unsuitability [2]. It is therefore critical to understand why greyhounds may fail to adequately perform the activity they were specifically bred to perform. This is a multi-faceted issue with many potential contributing factors, and little research has been conducted on it to date. Although it is likely that most behavioural wastage has taken place before greyhounds reach the track [1], a key component of understanding why greyhounds may fail to chase is in understanding their experience of race meets.

Greyhound races in Australia begin with a so-called stir-up, which is where greyhounds may watch the lure traverse the track usually twice while they are in a pen outside the track. Pre-stir-up occurs approximately ten minutes before the start of each greyhound race, and five minutes before the stir-up under Australian greyhound racing rules [3]. Pre-stir-up involves the collection of the dogs from their kennel and walking them to a grass area next to the track where they may eliminate, and fitting them with a racing rug and any additional pre-race preparations such as taping body parts for protection against injury. After witnessing the lure traverse the track, the greyhounds are walked to their starting boxes, loaded into the boxes, and then released from the boxes and chase the lure for a set distance. At the end of the race, a gate is swung across the track immediately behind the lure to stop the greyhounds from chasing it,. The lure itself draws away from the greyhounds and passes through a small flap in this gate, and the greyhounds are diverted into the catching pen alongside the track, where they are caught by handlers and then led from the track. The catching pen is unique to Australia, and, it is argued by industry participants [4], may be a source of frustration for greyhounds, as they are unable to capture the lure and there is rarely an object in the catching pen for them to interact with in lieu of capturing the lure. The consequences of frustration may be subsequent failure to chase or redirection of frustration onto nearby dogs, both of which attract penalties if they occur during races rather than in the catching pen. Furthermore, the risk of injury in the catching pen may be greater if greyhounds redirect frustration onto nearby conspecifics as race participants are decelerating at different rates.

Many factors influence performance of racing animals. Previous research on racehorses has shown that horses that finished as winners (top 20% of finishers) tended to be less aroused in the mounting yard immediately before the race than the losers (bottom 20% of finishers) [5], with arousal being determined by behavioural indicators. High arousal may lead to a reduction in fine motor control [6], and may also compromise judgement and cognitive processing [7]. These outcomes may manifest in racing greyhounds that show poor cornering or manoeuvring around other dogs, starting a race too fast or expending excessive energy in the stir-up and fatiguing early, or interfering with other dogs during the race. However, it is impossible to tell whether a dog is performing according to a typical pattern they employ on the racetrack, or if they are performing differently from usual, so it is necessary to seek indicators of sub-optimal arousal. We therefore employed the use of an infrared thermographic (IRT) camera to record the surface temperature of greyhound eyes before and after the race. IRT detects infrared radiation, providing a pictorial representation of surface temperature [8]. Typically, vascular perfusion of the extremities changes during stress responses, which include increased arousal. In animals, this can be detected by a change in the surface temperature of superficial, hair-coat-free, anatomical landmarks of an animal that are perfused by extensive capillary networks, such as the eye and inside the ears [9]. Due to parasympathetic activation, dogs may exhibit an increase in heart rate and peripheral vasodilation upon the onset of a stress response, resulting in increased metabolic heat production, and an increase in surface temperature, which is most easily detected on the surface of the eye [8]. Such increases in eye temperature can be detected by IRT, and has successfully assessed arousal in a variety of animals including mice [9], rabbits [10], horses [11–15] and dogs [8,16–18].

Heightened arousal prior to the race in greyhounds may be caused by distress related to the racetrack environment including kennelling. Anticipation may also heighten arousal levels [16,18,19], which may in turn be influenced by how long the dog has been kennelled for at the race meet, or how many days it has been since the dog last raced, or how experienced the dog is with the procedure at race-meets. A previous study on racing greyhounds revealed an increase in arousal in dogs that race as well as those that have merely watched racing [20], suggesting greyhound arousal increases with anticipation of an opportunity to race. Road transport over an hour in duration is regarded as distressing for livestock animals [21], and studies on air travel in dogs show it increases behavioural and physiological signs of distress [22, 23].

The current study aimed to determine possible effects of arousal and frustration on performance in racing greyhounds at race-meets. As well as obtaining IRT images of greyhounds before and after races, behavioural observations were collected of greyhounds during the stir-up immediately prior to racing, and in the catching pen at the conclusion of races to explore putative behavioural indicators of increased arousal before racing and signs of frustration in the catching pen associated with being thwarted in capturing the lure.

## Methods

The University of Sydney Animal Ethics Committee approved the current study (Approval number: 2016/1015). The owners/handlers of the greyhounds provided informed consent for the collection of infrared images.

### Location

The study was conducted at three greyhound racetracks in NSW over a period of 6 months. The tracks were Richmond and Wentworth Park in the Sydney metropolitan area in June and July 2017 respectively, and Gosford on the New South Wales Central Coast, approximately 80km north of Sydney, in October and November 2017. Data were collected from 3 race meets at Richmond, with 11 races per meet, 2 race meets at Wentworth Park with 10 races per meet, and 3 race meets at Gosford with 8, 10 and 11 races respectively.

Each track was configured differently (see supplemental material for diagrams). Minimum distances between features of the track and where on the grounds greyhounds were subject to potentially arousing stimuli were measured using the measurement tool in Google Earth Pro (Google Earth Pro version 7.3.2.5776, Google LLC 2019).

During the period of data collection, the Richmond racetrack was trialling a bungee teaser in the catching pen. Teasers consisted of two toys made of synthetic fur attached to one bungee line each that was in turn anchored at the back fence of the catching pen. To offer teasers, the track steward operating the catching pen gate walks onto the track with the teasers, stretching both bungee lines taut. The steward releases the teasers as the dogs approach the catching pen, providing a moving stimulus across the track and into the catching pen. The teasers come to rest in the sand trap of the catching pen and the dogs are able to interact with them. All greyhounds racing are muzzled, so interactions with the teaser are restricted. This system was in place for all race-meets where data were collected at Richmond.

### Dogs

A total of 525 greyhounds were recruited to this study over the 8 race meets at 3 racetracks. The races included were for both male and female greyhounds aged 1-6 years old, and dogs varied in experience, with their number of starts ranging 0-177. The dogs arrived at the racetrack in air-conditioned dog trailers, which is the current policy of Greyhound Racing NSW (GRNSW). Upon entry to the racetrack the dogs undergo vetting: an approximately 30-second veterinary physical examination to ensure quality of health. The dogs are then kennelled in an air-conditioned building where they remain until they are taken out at pre-stir-up, approximately 10-15 minutes before their race. Dogs were excluded from data collection if they had already been recorded by the current team of investigators at a prior race or race-meet.

### Physiological data collection

IRT data were collected twice from each dog during each race meet, with the first IR thermograph being taken during pre-stir-up. The second IR was taken 15 minutes after the race while the greyhound was kennelled. Post-race kennelling proceeds after greyhounds are hosed down and offered water to drink upon finishing the race.

IRT images were captured using a FLIR T640 Professional Thermal Imaging camera (T640, FLIR Systems Inc. Danderyd, Sweden) at an 80° or 100° angle and 1m distance from the dog. The FLIR ResearchIR Max software program was used to calculate the average eye temperature under the 1234 palette because it best exposed the circumference of the eye. Greyhound eye temperature was calculated by tracing the eye lids of the greyhounds using the Stats tool then using Statistics Viewer to calculate the mean and max temperature inside the traced area.

### Behavioural data collection

The behaviour of the dogs was recorded using one GoPro Hero3 Black Edition action camera (Manufacturer’s name and address) mounted onto the fence of the catching pen, and one hand-held Sony HD Handycam HDR-PJ760 video camera. The videos were analysed in slow motion in Windows Media Player 11 (Microsoft, Redmond, Washington, USA) (0.5x speed) and a count of each of the behaviours listed in the ethogram (Table 1 and 2) were recorded for each dog. Greyhounds whose trainers excluded them from the optional stir-up event were not analysed with an ethogram, and were recorded as being absent.

**Table 1:** Ethogram of all the behaviours potentially indicating arousal during stir-up in greyhounds.

| Behaviour | Description | Frequency |
| --- | --- | --- |
| Rising (r) | Unassisted rising onto hind legs without hind feet leaving the ground in vertical or lateral movement | Count |
| Owner-assisted rising (ro) | Owner lifting dog onto hind legs without hind feet leaving the ground in vertical or lateral movement | Count |
| Lunging (lu) | Lateral thrust forward, therefore pulling on leash | Count |
| Spinning (s) | Dog rotates laterally either clockwise or counter-clockwise for approximately a full revolution | Count |
| Jumping (j) | Both front and back feet leaving the ground so that a suspension phase occurs | Count |
| Barking (b) | Barking | Count |

**Table 2:** Ethogram of all the behaviours (with abbreviations) that potentially indicate negative emotional valence in greyhounds in the catching pen after a race

|  |  |  |
| --- | --- | --- |
| Grabbing the teaser (gt) | Teeth contact the teaser but are released before trainers contact the dog | Yes/no |
| Changing directions (cd) | An approximate 180° change in direction while in motion. | Count |
| Jostling (jo) | Dog's muzzle makes physical contact with another dog with sufficient force to affect the receiving dog | Count |
| Attention directed to lure gate on racetrack (fl) | Dog orientating body position and interest towards the lure gate, or attention to the lure as it slows down around the track | Yes/no |
| Holding teaser (ht) | Teaser is grabbed and not released by the greyhound | Yes/no |

The ethograms were informed in part by Travain et al 2015 [8], who used an ethogram to estimate distress in a group of 14 dogs, with behaviours being considered indicative of distress when they were accompanied by a significant increase in eye temperature (detected by IRT) [8]. No racing-specific ethogram for dogs has been developed before, so several behaviours were added to the ethograms that were considered good candidates for detecting high arousal, frustration, or fixation on the lure.

### Questionnaire data collection

The questionnaire for trainers consisted of 4 questions that were used to identify any indirect factors that may influence affective state:

1. How long did it take you to get to the racetrack (minutes)?
2. How many times has your greyhound raced (starts)?
3. How long since the greyhound’s last race (days)?
4. How old is your greyhound (years, rounding to the nearest half year.)?

The data from this questionnaire were compiled along with the greyhounds’ start box number, the date of the race, track, race distance, performance (placing), time of the race meet, and ambient temperature at the time of the dog’s race. Ambient temperature was collected from records available from Time And Date AS, which purchases weather information from customweather.com [24]. The records are available on an hourly basis for the Richmond, Sydney city, and Gosford localities.

### Statistical analysis

All statistical analyses were performed in RStudio (version 1.1.383, desktop macOS, RStudio Inc., Boston, Massachusetts, USA). Behaviours were pooled into three categories to address some low counts. These categories were: “Aroused_S for behaviours” indicating arousal during stir-up, “Unresolved” for behaviours in the catching pen that may indicate the greyhound was still fixated on the unattainable lure or expressing frustration, and “Teaser” for behaviours in the catching pen at Richmond that were related to interacting with the teasers on bungee lines. The frequency of behaviour recordings both in the catching pen and in the stir-up were rarely more than 5 counts. The only exception was barking, which is energetically a much less costly behaviour and is also much quicker to perform than other behaviours in the ethograms. All behaviours were scaled using the max-min method to a scale of 0-5 counts to avoid the inflation of results in dogs prone to vocalisation. Counts for pooled behaviours were then rounded to the nearest whole number to allow for a negative binomial model to be fitted. This step was relevant only for Aroused_S behaviours, as behaviours in other categories did not need to be scaled.

An ordinal linear regression model was used to determine factors influencing performance. Generalised linear models with a quasi-poisson distribution due to over-dispersion in count data were used from the lme4 package using the glm function in RStudio to determine factors that have a significant effect on Aroused_S behaviours and Mean ET Before races and Mean ET After races. The final models were built using the stepwise method and the AIC number to determine the model of best fit.

Pearson’s Correlation tests using the cor function were performed on factors that were not included in models or for which models were difficult to resolve.

## Results

### Tracks

Track configuration in terms of where the kennel block, stir-up yard and catching pen were located in relation to the track differed between tracks, as summarised in Table 3.

**Table 3.** Distances in metres between features of each race track in the study.

| Track | Kennels to stir-up (m) | Kennels to track (m) | Stir-up to track (m) | Catching pen to kennels (m) | Catching pen to stir-up (m) |
| --- | --- | --- | --- | --- | --- |
| Richmond | 5 | 34 | 18 | 140 | 123 |
| Wentworth Park | 1 | 8 | 1 | 5 | 2 |
| Gosford | 17 | 28 | 3 | 170 | 150 |

### Performance

The final ordinal linear regression model on greyhound performance included the factors mean eye temperature before the race (MeanETBefore) mean eye temperature after the race (MeanETAfter), dog age (Age), start box number (Box), number of dogs in the race (Field), the number of days since the dog last raced (Days_last_race) and sex of the dog (Sex). It also included an interaction between Sex and Days_last_race. Increasing MeanETAfter had a negative effect on performance, as shown in Figure 1 (n=290, Effect = −0.171, s.e. = 0.073, p-value = 0.019) and increasing age had a negative effect on performance, shown in Figure 2 (n=290, Effect = −0.395, s.e. = 0.136, p-value = 0.004). On the whole, male dogs performed better than female dogs (n=290, Effect = 0.752, s.e. = 0.257, p-value = 0.003), but they performed worse with increasing number of days since they were last raced, as shown in Figure 3 (n=290, Effect = −0.022, s.e. = 0.010, p-value = 0.023). This is demonstrated further in Figure 4, which shows the predicted placings of male dogs given weeks since last raced when all other factors are held constant. The probabilities associated with those predictions are shown on the y-axis. These figures were obtained from the same ordinal model run on a subset of the original data containing only male dogs.

**Figure 1:**
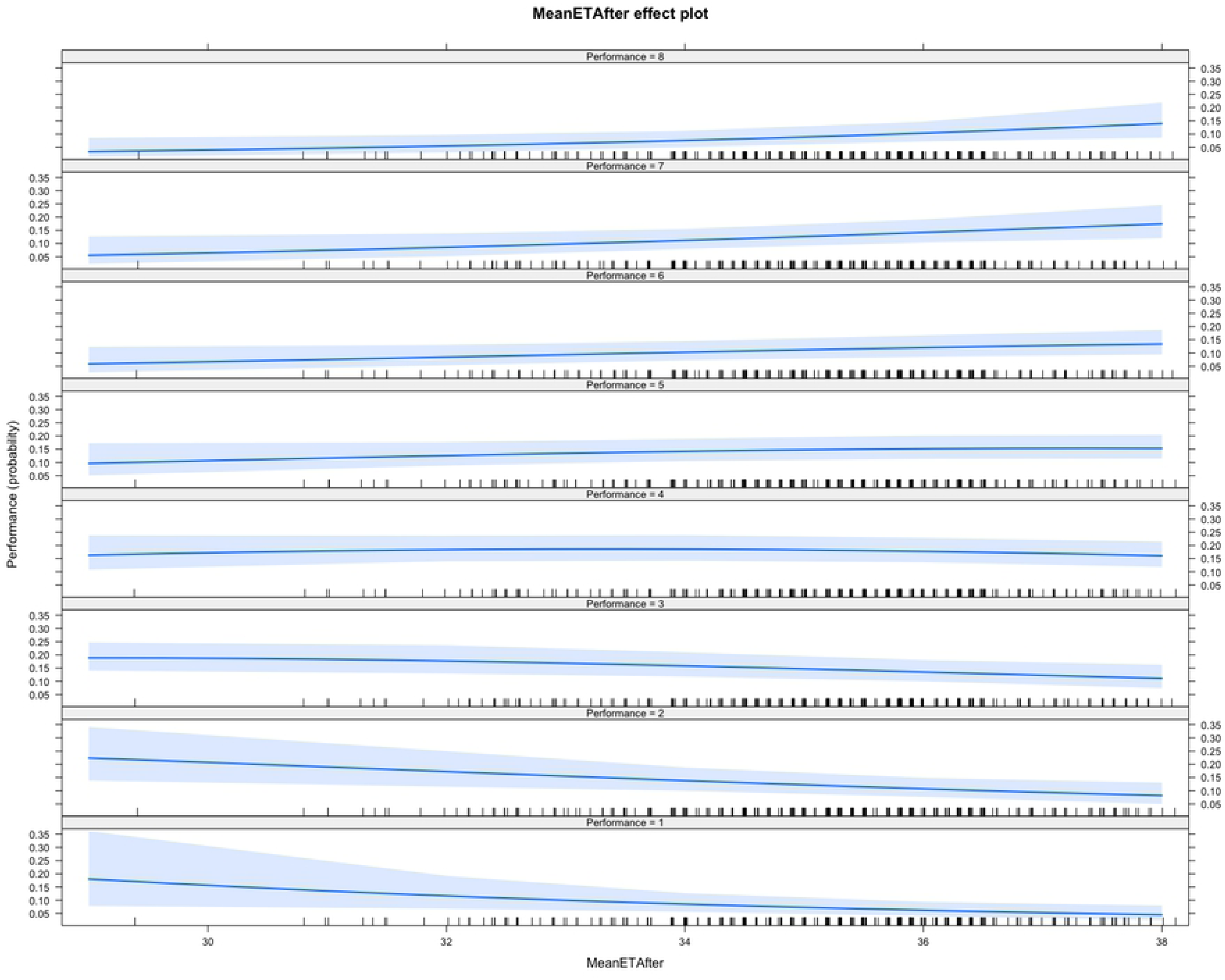
The probabilities of different mean eye temperatures 15-minutes after racing for placing 1^st^ through to 8^th^ where races include 5-8 dogs. Mean eye temperatures (Celsius) are shown on the x-axis. Probability of mean eye temperature given placing (1^st^-8^th^) is shown on y-axis. 95% confidence intervals are shown in shading. There is a higher probability of higher mean eye temperature in dogs with poorer placings.

**Figure 2.**
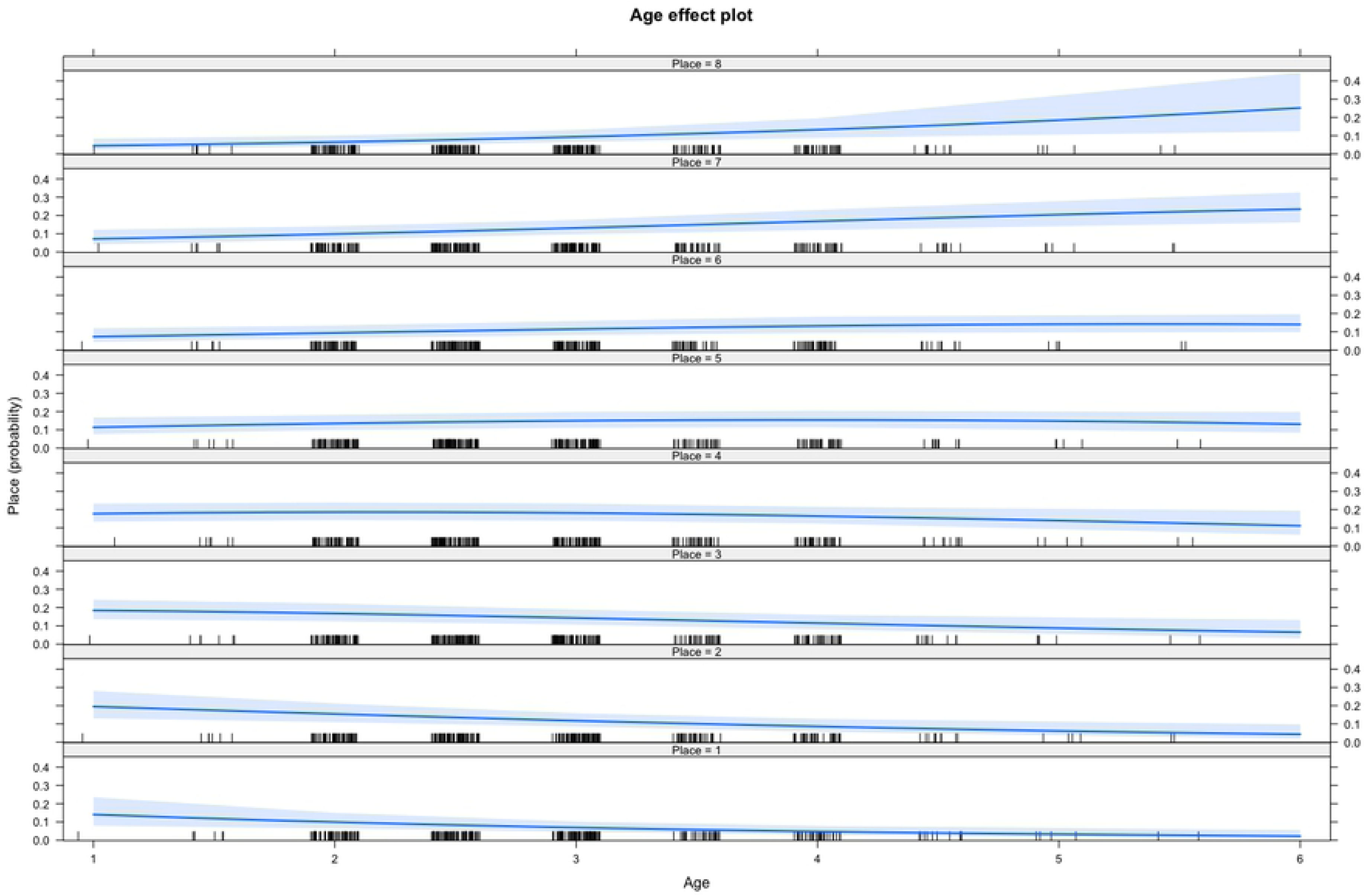
The effect of greyhound age (x-axis) on the probability (y-axis) of placing (Place 1-8). 95% confidence intervals shown with shading. Younger dogs were more likely to place favourably (Place 1-4) than older dogs.

**Figure 3.**
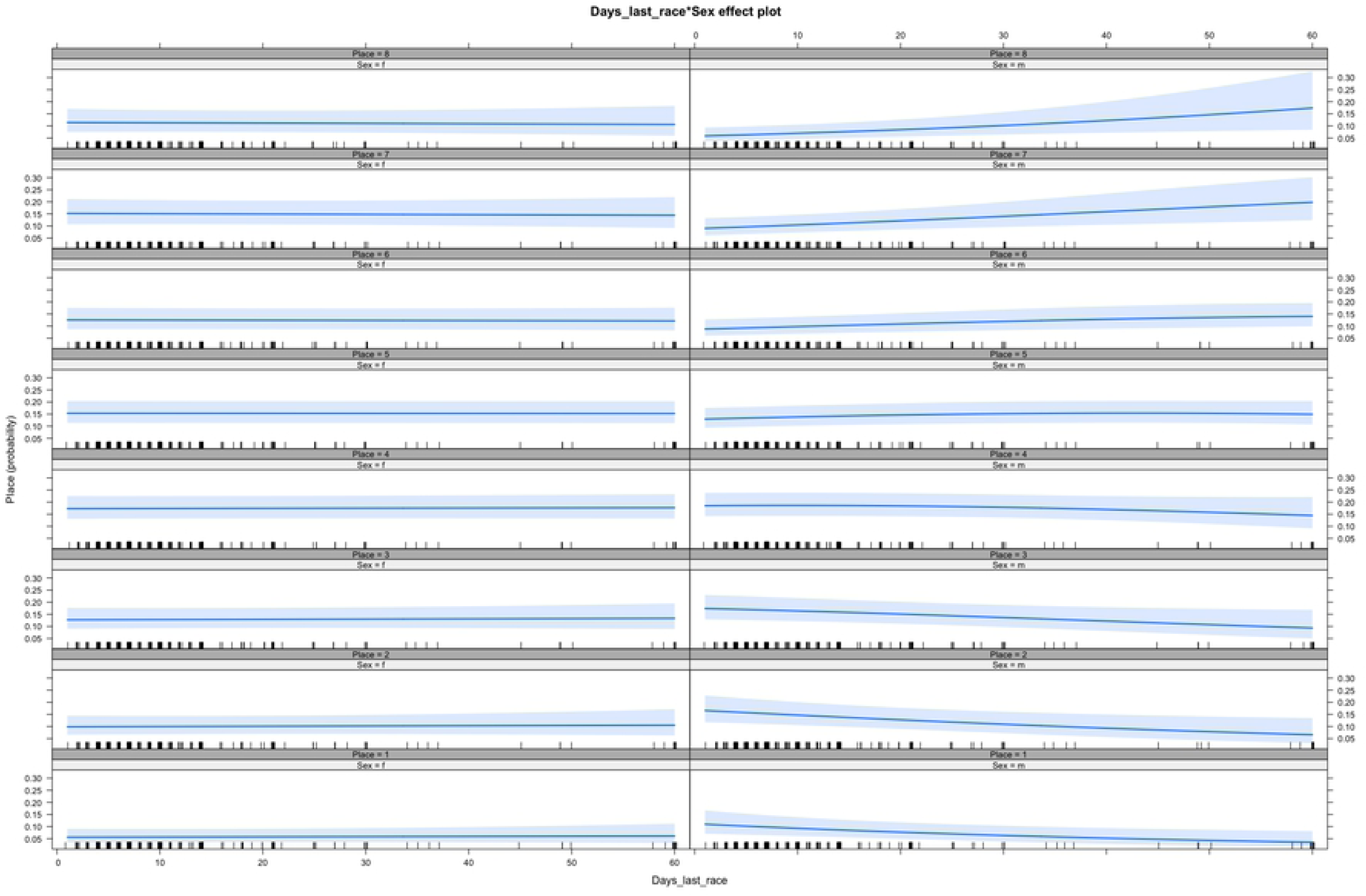
The effects of days since last raced (x-axis) on the probability of placing (Place 1^st^-8^th^) for females (left column) and males (right column). There was an interaction between sex and days since last raced, with males more likely to place poorly as intervals since last racing increased. 95% confidence intervals shown in shading.

**Figure 4.**
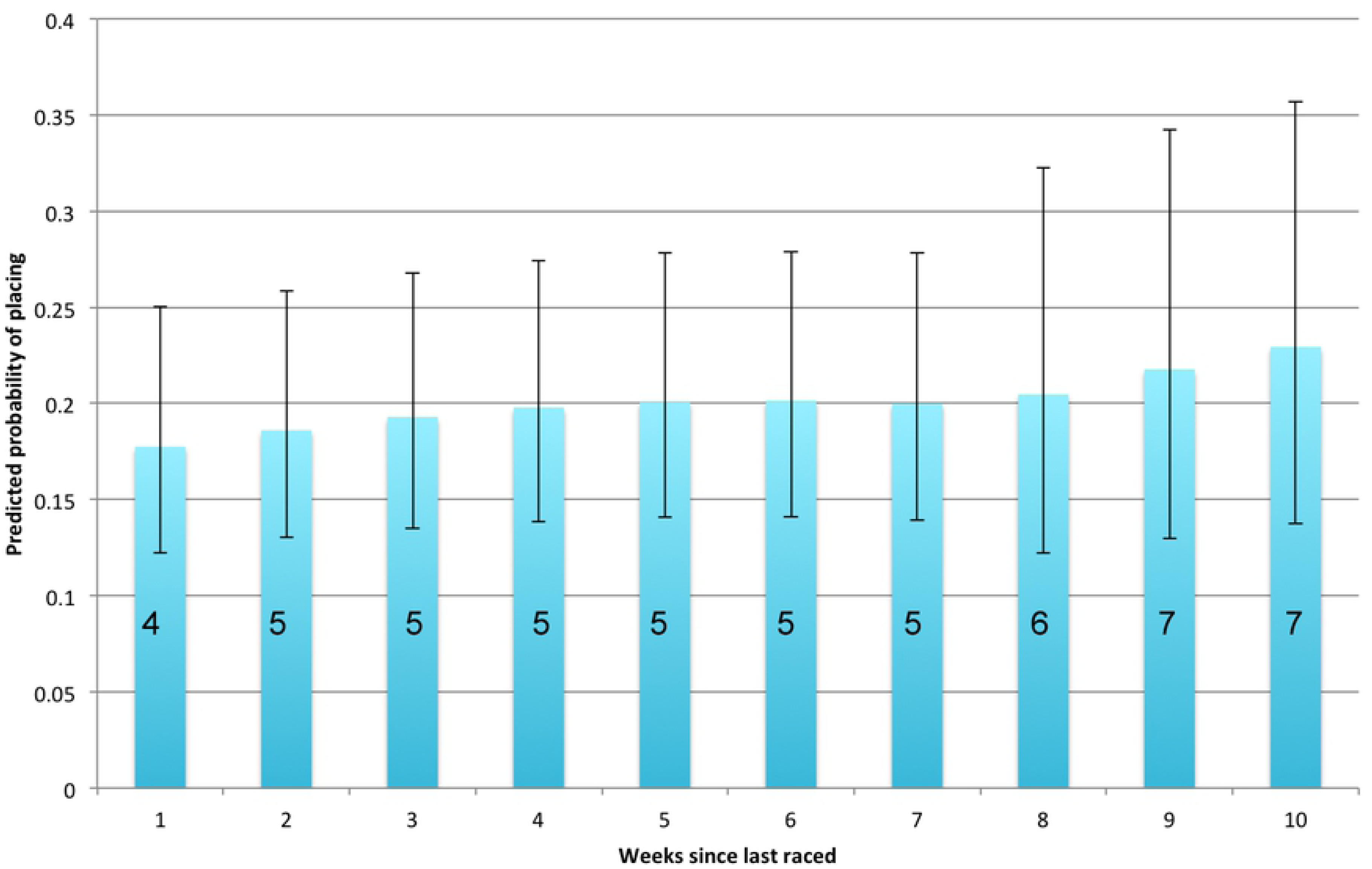
Predicted placings of male dogs 1-10 weeks after since last racing. Places with highest probability when all other factors in the ordinal linear regression model are held at their mean are shown on the bars. Error bars show the upper limits and lower limits of predicted probabilities associated with the placings. Thus, there is a predicted loss of 3 places between racing male dogs a week after their last race compared to 10 weeks after their last race.

Box 1 showed the strongest association with good performance while, in comparison, Boxes 4, 5 and 7 had a significantly negative effect on performance (see Table 4 for figures). Figure 5 shows the probability of placings from each starting box. The other factors did not have a statistically significant effect on performance, but their presence improved the model according to the AIC. The results of this model are shown in Table 3.

**Figure 5.**
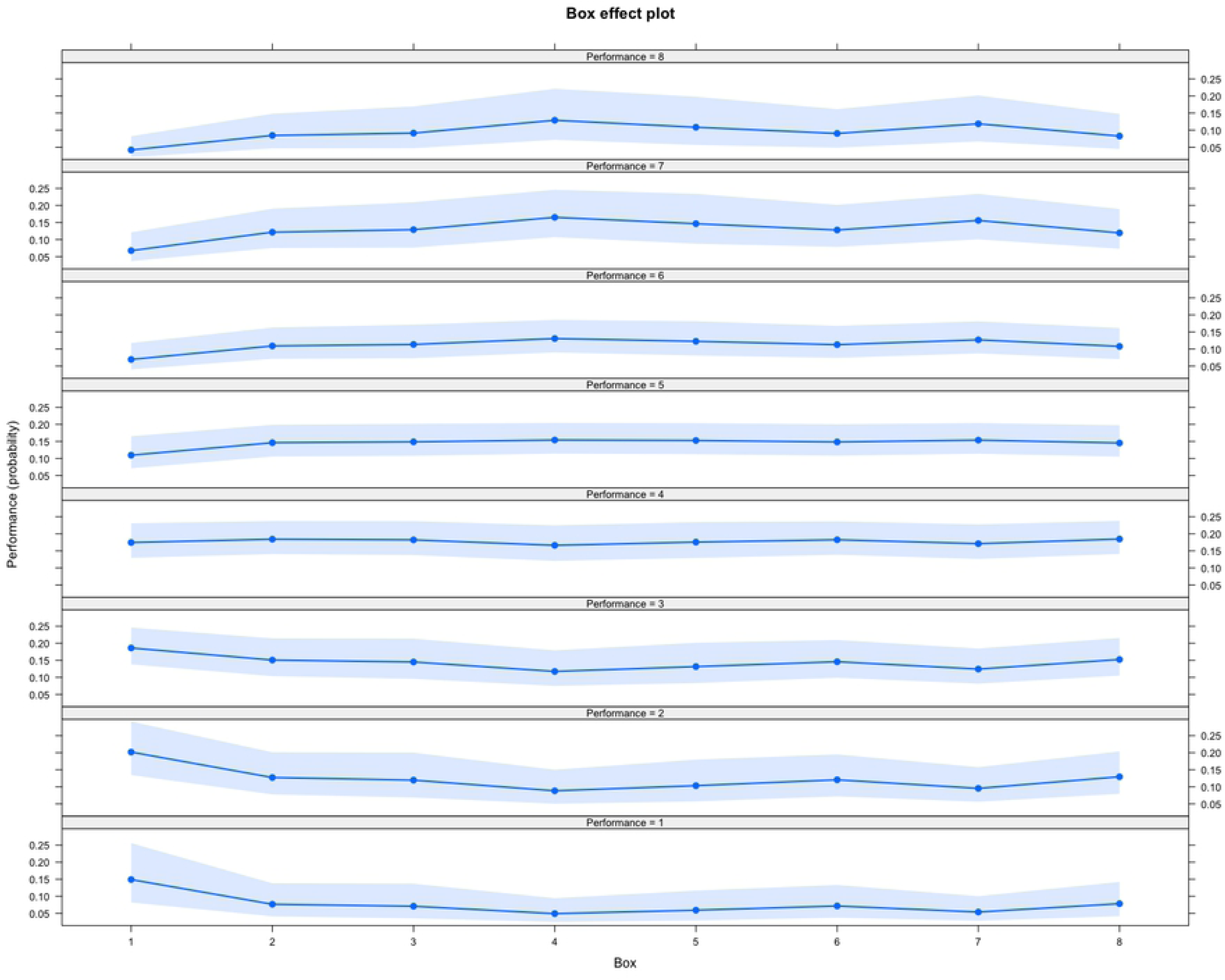
The effects of the starting box on the probability of placing 1^st^-8^th^ (Performance). Dogs have a higher probability of placing first or second if they start from Box 1. Boxes 4 and 5 are associated with an elevated probability of placing 7th or 8th. 95% confidence intervals shown in shading. Box is on the x-axis and probability given placing (Performance) is on the y-axis.

**Figure 6:**
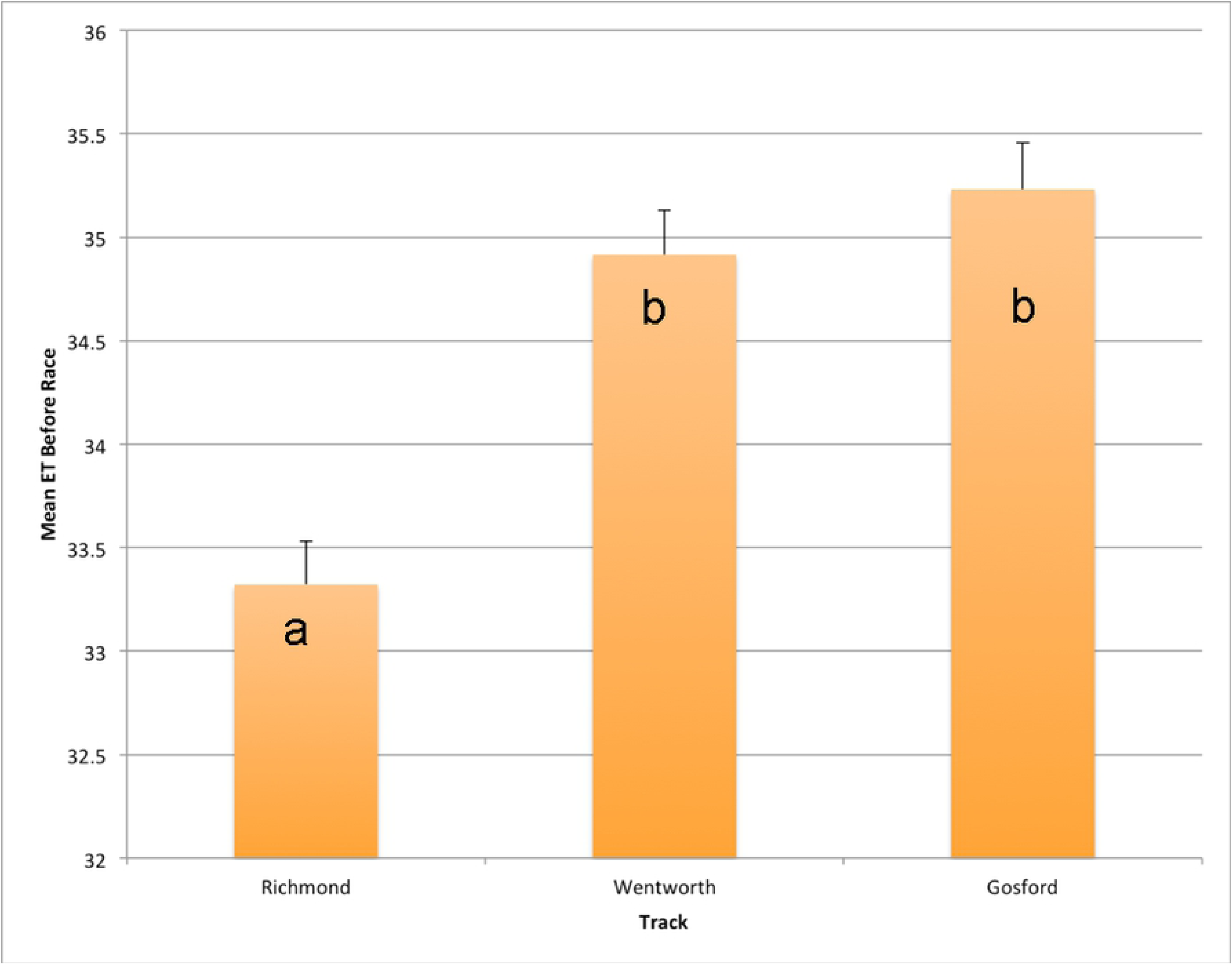
Fitted Mean ET Before races at different tracks. MeanETBefore was much lower at Richmond than the other two tracks, suggesting dogs are overall calmer at Richmond racetrack than Wentworth Park or Gosford. Superscripts are assigned to values that are significantly different from one another.

**Table 4.**
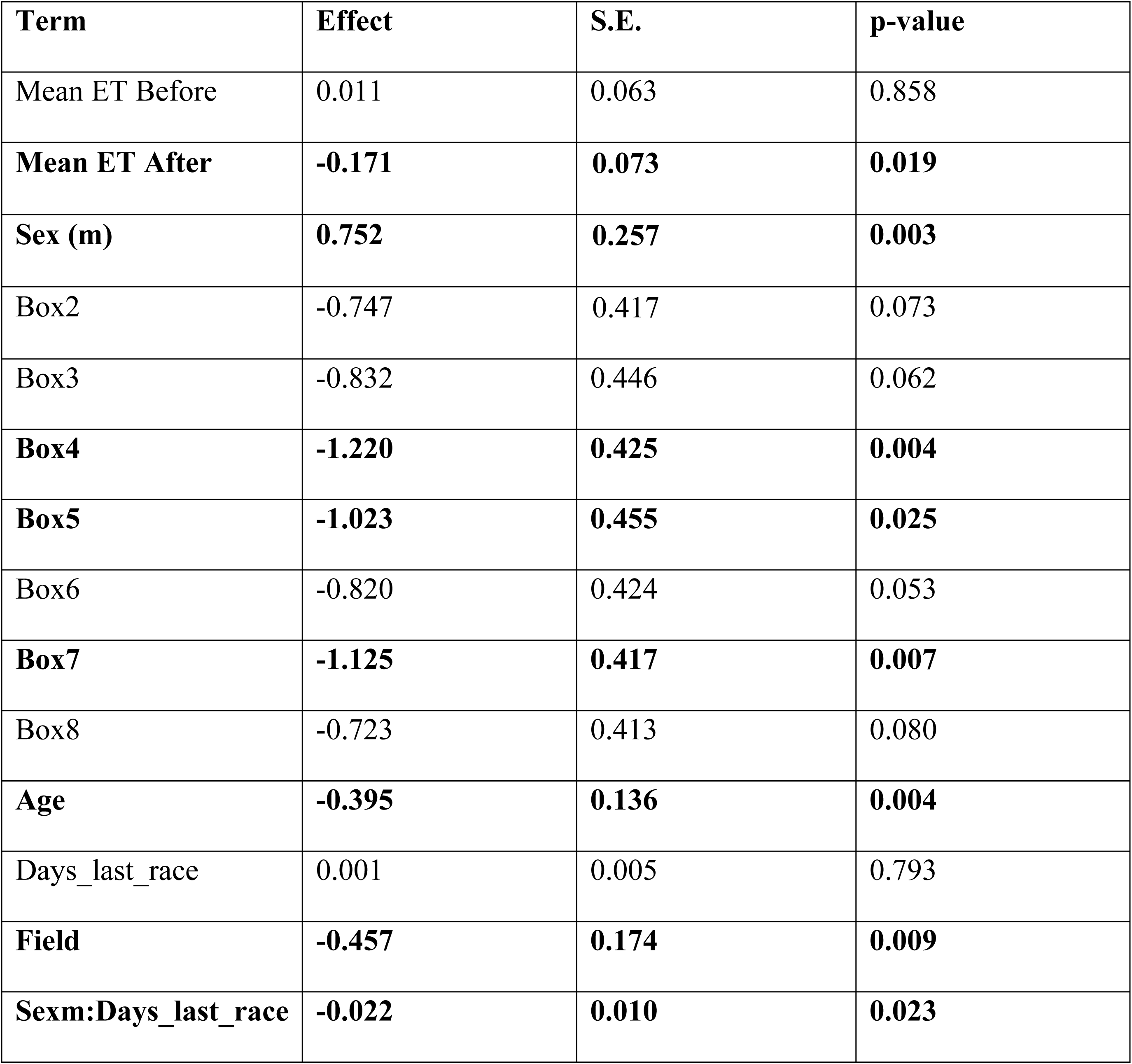
Summary of an ordinal linear regression model on performance in racing greyhounds. Factors with a significant effect on performance appear in bold. Mean eye temperature 15-minutes after the race, Start Boxes 4, 5 and 7, increasing age, and days since last raced for males only had a negative impact on performance. Male dogs performed better than female dogs. The number of dogs in each race (Field) was included in the model to account for the possible effects of there being fewer dogs in the race.

### Aroused behaviours

The negative binomial model on the frequency of aroused behaviour was constructed in the same manner as the ordinal model. This model included mean eye temperature before and after the race, race distance (Distance), racetrack (Track), sex, and Days_ last_race. A summary of these results is shown in Table 5. Increasing race distance had a negative effect on the frequency of aroused behaviours in the stir-up (n = 290, Effect = −0.004, s.e. = 0.002, p-value = 0.031), and the race being held at Wentworth Park had a positive effect on the frequency of aroused behaviours in the stir-up (compared to Gosford) (n = 290, Effect = 1.255, s.e. = 0.380, p-value = 0.001). There was a non-significant trend for increasing number of days since last raced to be associated with reduced Aroused_S behaviour counts.

**Table 5:**
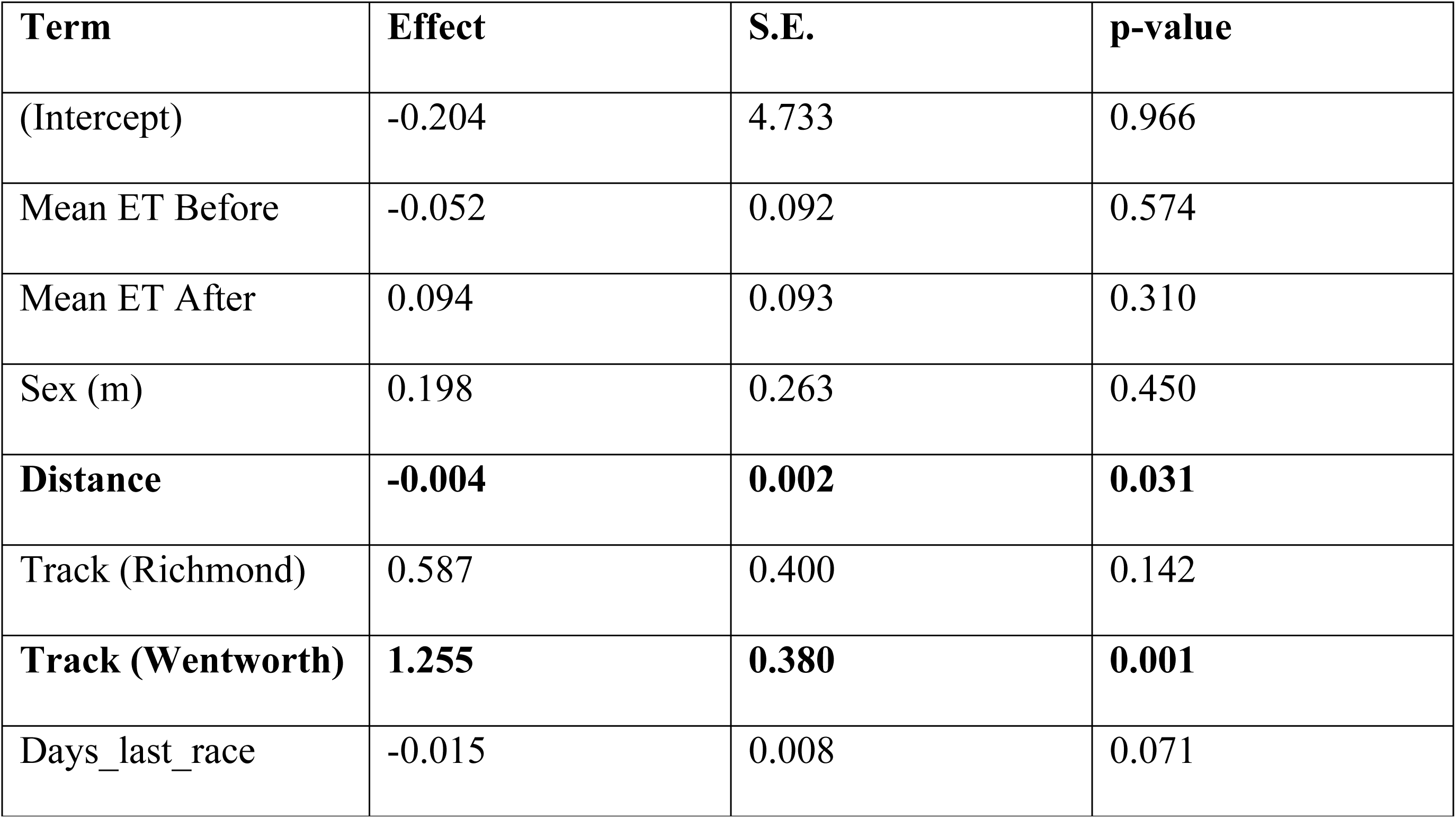
Summary of a negative binomial linear regression model of aroused behavior during stir-up (immediately before racing). Statistically significant results are shown in bold. Increasing race distance had a negative effect on the frequency of aroused behaviours. More aroused behaviours were observed at the Wentworth Park track than at the Gosford track.

| Term | Effect | S.E. | p-value |
| --- | --- | --- | --- |
| (Intercept) | -0.204 | 4.733 | 0.966 |
| Mean ET Before | -0.052 | 0.092 | 0.574 |
| Mean ET After | 0.094 | 0.093 | 0.310 |
| Sex (m) | 0.198 | 0.263 | 0.450 |
| **Distance** | **-0.004** | **0.002** | **0.031** |
| Track (Richmond) | 0.587 | 0.400 | 0.142 |
| **Track (Wentworth)** | **1.255** | **0.380** | **0.001** |
| Days_last_race | -0.015 | 0.008 | 0.071 |

### Mean eye temperature

The generalised linear model for Mean ET Before races contained the terms Track, Sex, Race and Aroused_S, and a summary of the model is shown in Table 6. Track had a powerful effect on Mean ET Before, with both Gosford and Wentworth having a strong, positive effect, as shown in Figure 5 (n = 442, Effect = 1.910, s.e. = 0.152, p-value = 0.001; Effect = 1.595, s.e. = 0.159, p < 0.001 for Gosford and Wentworth respectively). Increasing race number had a strong, positive effect on Mean ET Before (n = 442, Effect = 0.103, s.e. = 0.022, p-value < 0.001), s shown in Figure 7. Males had lower Mean ET Before, and Aroused behaviours in stir-up had a positive effect on Mean ET Before, but the effect of both factors was very small and not statistically significant. A generalised linear model for Mean ET After races contained the terms Race, Temperature, Track, Distance, Sex, Aroused_S and Unresolved, and revealed that statistically significant predictors of Mean ET After were ambient temperature (n = 310, Effect = 0.149, s.e. = 0.032, p-value < 0.001) and race number had a positive effect (n = 310, Effect = 0.071, s.e = 0.027, p = 0.010) (Table 7). A scatter plot showing the relationship between ambient temperature and Mean Eye Temperature Before the race is shown in Figure 8. Mean Eye Temperature After the race was, as expected, influenced by ambient temperature, and the relationship is shown in Figure 9.

**Figure 7:**
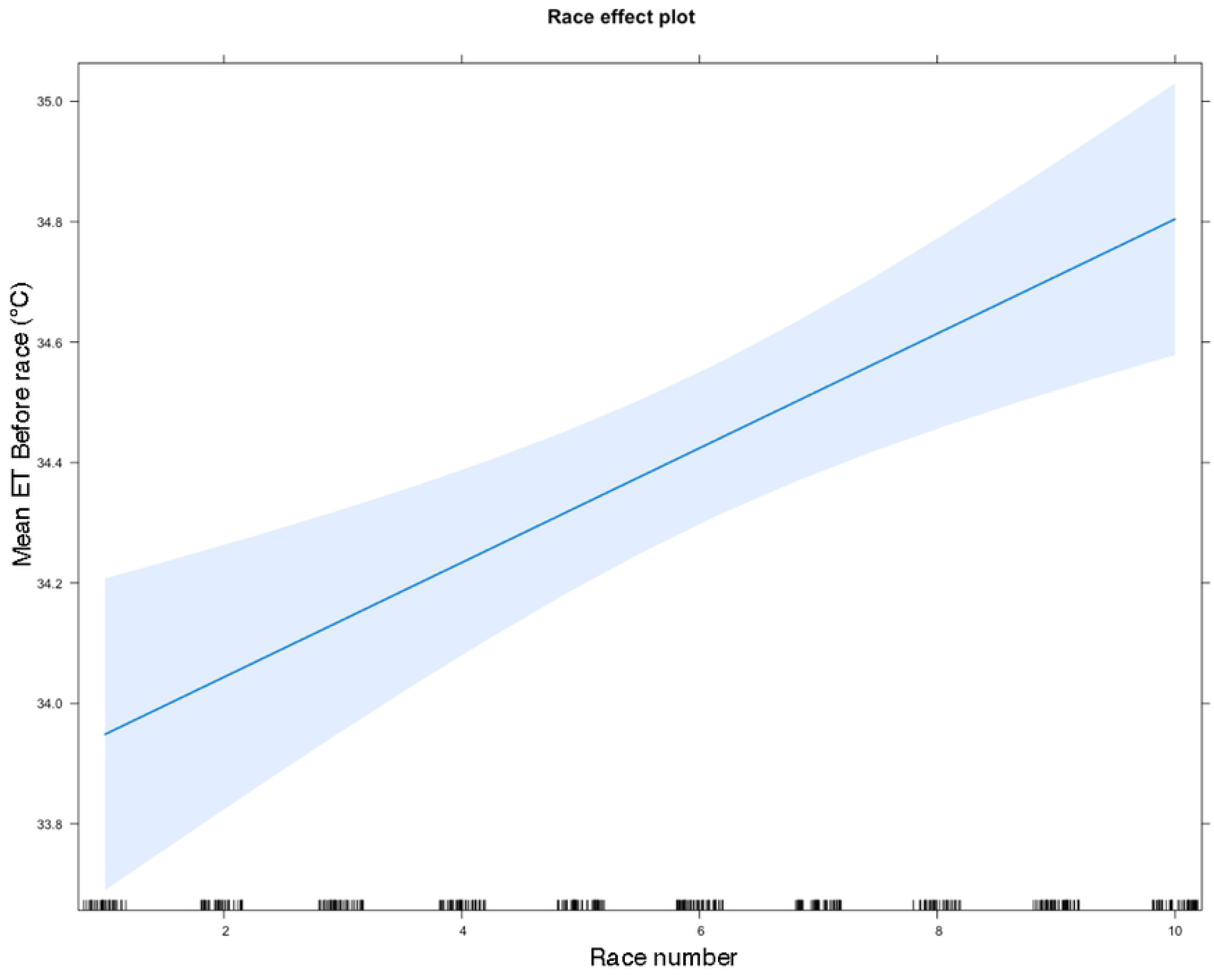
Predicted Mean ET Before races depending on race number. Mean ET Before races increases as race number increases, indicating greyhounds become increasingly aroused as the race meet progresses. 95% confidence intervals shown with shading.

**Figure 8:**
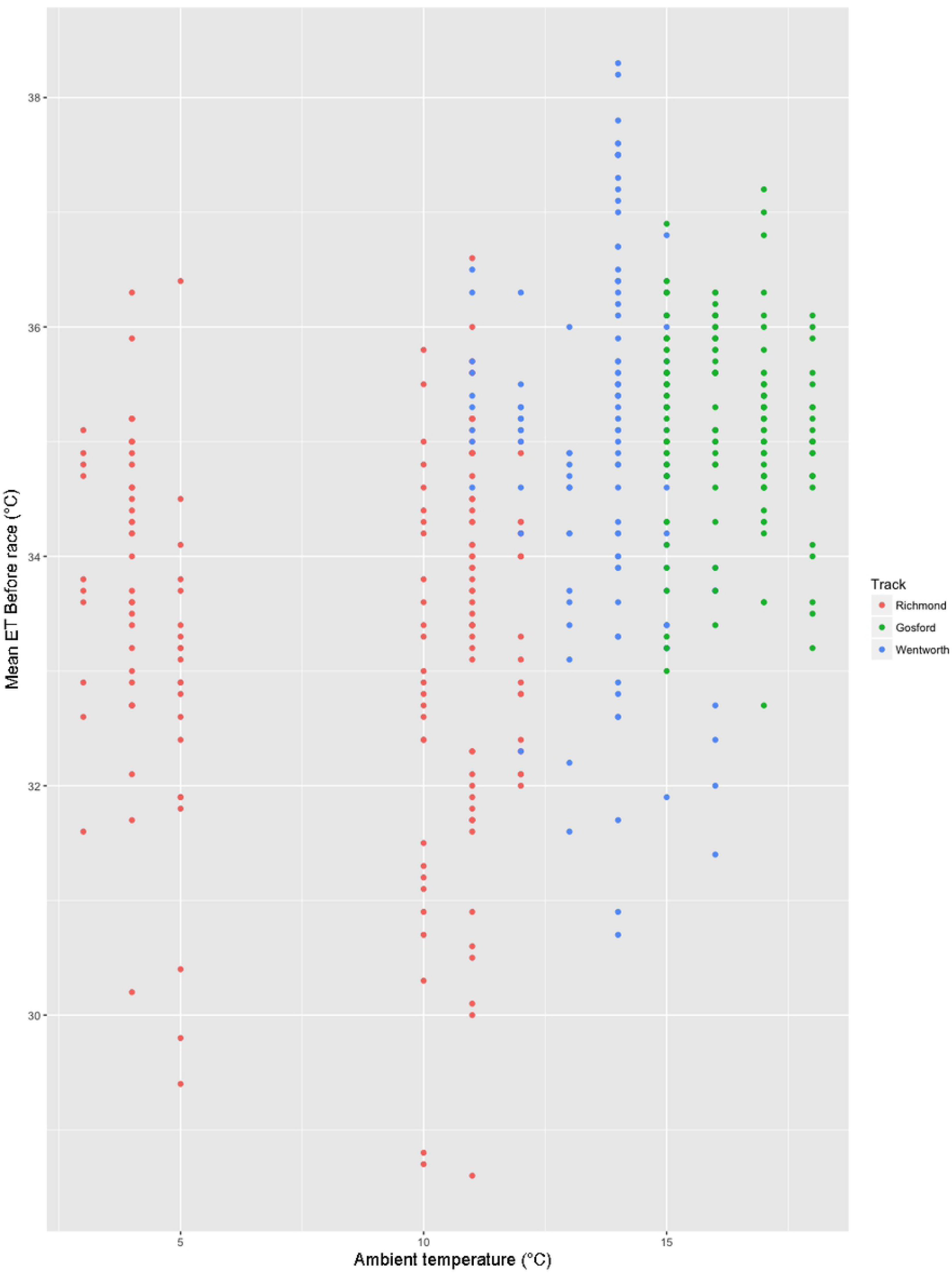
Scatter plot showing relationship between Mean Eye Temperature Before the race and ambient temperature for each track. Ambient temperature was not included in the model to predict Mean Eye Temperature Before because its addition to the model worsened the fit of the model according to the AIC.

**Figure 9.**
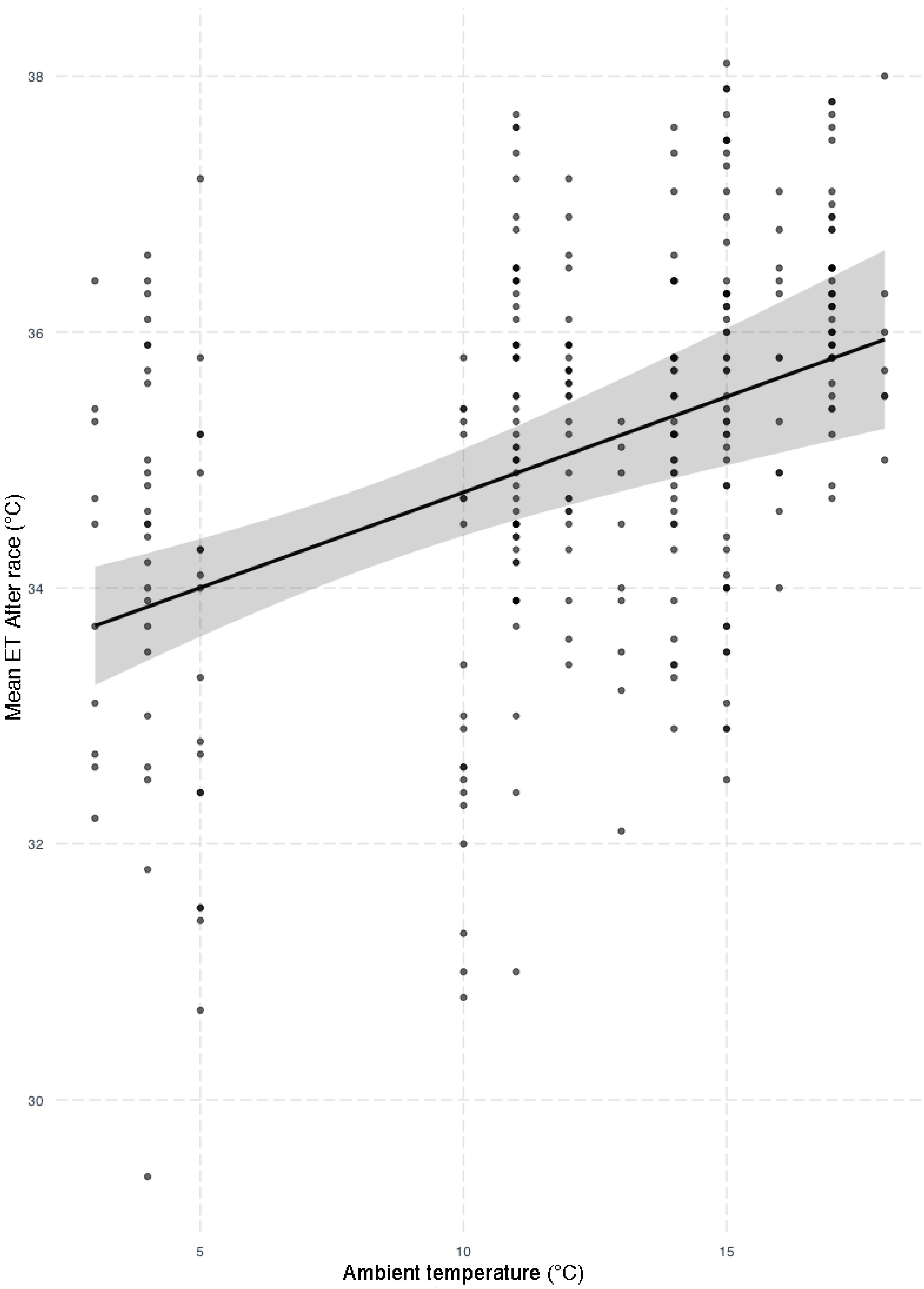
Predicted response of Mean Eye Temperature After the race to ambient temperature across all tracks. Ambient temperature significantly influenced Mean Eye Temperature After the race, but not greyhound performance.

**Table 6:** Generalised linear model summary for Mean ET Before races. Statistically significant effects appear in bold. Track and increasing race number both have significant, positive effects on Mean ET Before races.

| Term | Effect | S.E. | p-value |
| --- | --- | --- | --- |
| (Intercept) | 32.676 | 0.185 | $< 0.001$ |
| **Race** | **0.103** | **0.002** | **$< 0.001$** |
| **Track (Gosford)** | **1.910** | **0.152** | **$< 0.001$** |
| **Track (Wentworth)** | **1.595** | **0.159** | **$< 0.001$** |
| Sexm | -0.007 | 0.130 | 0.959 |
| Aroused_S | 0.005 | 0.034 | 0.872 |

**Table 7:** Generalised linear model summary for Mean ET After races. Statistically significant effects are in bold. Ambient temperature (Celsius) and increasing race number both have significant, positive effects on Mean ET After races.

| Term | Estimate | Std. Error | P-value |
| --- | --- | --- | --- |
| (Intercept) | 33.416 | 0.704 | <0.001 |
| **Race** | **0.071** | **0.027** | **0.01** |
| Distance | -0.001 | 0.001 | 0.258 |
| Sex (m) | -0.014 | 0.157 | 0.927 |
| Track (Gosford) | 0.29 | 0.346 | 0.402 |
| Track (Wentworth) | -0.05 | 0.325 | 0.878 |
| Aroused | 0.04 | 0.041 | 0.328 |
| Unresolved | 0.055 | 0.136 | 0.688 |
| **Temperature** | **0.159** | **0.032** | **<0.001** |

There was a significant, negative correlation between Teaser-related behaviour and Mean ET Before at Richmond track where the teasers were available in the catching pen (n = 166, cor = −0.140, df = 446, p-value = 0.003). We were unable to resolve a model for the frequency of Unresolved behaviours in the catching pen or Teaser-related behaviours, which may be due to low count data combined with multiple factors having a small influence on these behaviours. However, the frequency of Unresolved behaviours in the catching pen at Richmond racetrack was dramatically lower (17.1% of starters) compared to Wentworth Park (77.1% of starters) and Gosford (96% of starters) racetracks.

## Discussion

This study focused on the behaviour and arousal of racing greyhounds during race-meets, and the effect of these factors on performance.

### Eye temperature

Dogs with higher mean eye temperatures after the race were more likely to place in the back half of the field. Eye surface temperature, as measured by IRT, has been shown to drop in companion dogs in response to separation from their owners [18], but to increase in response to anticipation of a reward [16], owner return after separation [18], or being subjected to a veterinary examination [8]. Previous studies on cattle, horses and chickens have shown a drop in eye temperature when the animal appears to be in pain, is under restraint, or has been startled [11,25,26], suggesting that this is consistent with an acute stress response dominated by activation of the sympathetic nervous system, whereas an increase in ET may be consistent with a stress response dominated by parasympathetic activation [8]. Anticipation of reward and veterinary examinations may represent a stress response dominated by the activation of the parasympathetic nervous system instead of the sympathetic nervous system, which may represent a more general and less acute stress response. This outcome would be consistent with the general, non-specific stress of a race meet, kennelling, travel, and environmental noise. However, if generalised stress and/or anxiety related to the race meet environment were influencing performance, this should have been detectable through a significant increase in mean eye temperatures before the race as performance decreased. That pattern existed in the current study, but was not statistically significant.

. Sampling in the kennels prior to racing would likely provide a better comparison of before and after race as the act of taking the dogs out of the kennels may elevate their aorusal. However, this was not permitted by the racing officials. Under the rules of racing [3], greyhounds must remain in the kennels for 15 minutes before they can be taken home, so the IR thermograph needed to be taken at this 15-minute juncture to allow the greyhounds to cool down so as to minimise the effects of body temperature on eye surface temperature, while still obtaining thermographs before the greyhounds were removed from their kennels.

### Mean ET After races

Higher mean eye temperatures after the race was associated with poorer performance, but it is difficult to separate the effects of physical exertion in racing from the effects of emotional state before and during the races. Studies have found that ET increases in response to physical exercise in dogs and horses [12,13,17,27], but the form of exercise in these studies was prolonged rather than the short intensity of sprint races in the current study. Breed differences in ET in response to physical exercise has been reported in horses [12] and dogs [17]. This is of particular interest in horses, as Bartolomé et al. [12] conducted their study on horses at a show-jumping competition, and suggested that some horses prone to higher emotional reactivity showed a stronger change in ET after performing. The horse breeds studied were more similar in size than the Labrador retrievers versus the beagles used in Zanghi et al.’s study [17], and the show-jumping competition more closely resembles the current racemeets than the free play-bouts studied by Zanghi et al. [17]. Thus, it is possible that the current mean eye temperatures after racing are indicative of a stronger stress response to racing, and the disruptive effects of over-arousal on performance.

An alternative possible reason for the inverse relationship between mean eye temperatures after the race and performance is that increased mean eye temperature after racing is more indicative of higher core body temperature than directly of emotional state. A previous study suggested it takes at least 30-minutes for dogs’ core body temperature to return to baseline after 30-minutes of exercise [17]. In the current study, it was not possible to collect IRT images reliably more than 15-minutes after racing at the racetracks due to owners removing them from kennels and transporting them home. As such, the negative relationship between performance and observed mean eye temperatures after the race may be a result of dogs that perform poorly having to expend more effort to compete in the race than dogs that perform well, and thus having higher core body temperatures and mean eye temperatures after the race.

### Ambient temperature

Ambient temperature was a significant predictor of mean eye temperatures after the race, but not mean eye temperatures before the race. This variable was not included in the performance model because it compromised model fit. Greyhounds are held in temperature-controlled kennels before their race, but extreme ambient temperatures may influence IRT images after races, and this may need to be treated with caution if the industry elects to use IRT in future. Further research into eye temperatures after highly arousing activities and various intensities of physical exercise will reveal the utility and limitations of IR images in assessing emotional states in dogs of various breeds and levels of fitness.

### Race number

There was a statistically significant, positive effect of race number on mean eye temperatures before the races, suggesting that greyhounds at the race-meet grew increasingly aroused as the race-meet progressed. All greyhounds racing must be kennelled 30 minutes before the first race. They are undisturbed in the time between kennelling closing and the first race, but once races are underway, a steady stream of trainers enter the kennels to collect dogs and return dogs that have just raced. The kennelled dogs are therefore exposed to ongoing disturbance, and also likely hear arousing auditory stimuli from the course, such as the sound of the lure moving on the track. Whether the kennels themselves and the noise of unfamiliar neighbouring dogs within them are a source of distress for greyhounds at race-meets or whether arousal in kennelled greyhounds rises with each anthropogenic disturbance is unclear and beyond the scope of the study, but the increase in mean eye temperatures before the races over the course of the race-meet suggests a continuous rise in arousal. This is unlikely to be related to the intensity of competition, as prize money does not routinely increase with each race. Greyhounds cannot know when they will be taken out of their kennel for racing, so may anticipate this occurring every time they are disturbed by the movements of trainers and dogs, leading to increasing frustration and aroused anticipation when they are not removed.

### Aroused behaviours in stir-up

A previous study found that horses displaying behaviour indicative of higher arousal immediately prior to racing performed more poorly than horses that appeared calmer [5]. We did not find such a relationship either between mean eye temperatures before the race or the frequency of aroused behaviours in the stir-up and performance in the racing greyhounds in this study. Indeed, there was no relationship among the behaviours in the current ethogram thought to indicate emotional arousal, mean eye temperature before or after the race, and performance. Aroused behaviours in stir-up were best explained by race distance and race venue, with fewer behaviours indicative of arousal being observed before longer race distances, and more of such behaviours at Wentworth Park racetrack than the other tracks. These results call into question both the validity of ethograms in assessing arousal at race meetings, and whether greyhounds encounter elevated arousal at least several minutes before the mandatory stir-up. It was not feasible to collect IRT images of greyhounds during or immediately following stir-up due to the pressing schedules of race meetings, so it is possible many greyhounds were not undergoing heightened arousal when the pre-race IRT images were being taken. Alternatively, the behaviour of dogs during stir-up may be poor indicators of arousal. Dogs that express their arousal overtly with easily detectable behaviours may be no more aroused than dogs that are passive during stir-up. Any difference in behaviours may not directly reflect a difference in arousal level.

### Age and experience

Younger dogs were more likely to place in the front half of the field than older dogs. This may be because younger dogs are less likely to be burdened with the effects of previous injuries or general degenerative changes that may occur with age. In racehorses, studies have found that the risk of injury increases with age [28, 29] and that racing speeds peak [30] or plateau [31]at approximately 4.5 years of age.

### Start box

The box from which dogs started also had a significant impact on their likelihood of placing favourably, an outcome which is freely acknowledged by track administrators, for Wentworth Park at least [32]. Box 1 appears to offer an advantage, as noted by The Greyhound Recorder [33] and, in the current study, Boxes 4, 5 and 7 conferred a significant disadvantage when compared to Box 1. Greyhounds that prefer to run close to the rail are likely to perform better regardless of starting box because they must cover less ground over the course of the race than greyhounds that prefer to run on the outside of the pack. As such, greyhounds that prefer to run close to the rail and that also start from Box 1, 2 or 3 are likely to cover less ground than greyhounds with this preference that start from boxes farther from the rail. This issue may be best addressed by adopting track safety recommendations for a lure system that places the lure closer to the centre of the track [34].

### Sex of dog

Male greyhounds in the current study were significantly more likely to place favourably than females. However, this was complicated by an interaction between sex and days since last raced. Whereas females showed no clear pattern in their performance regardless of how long it had been since they were last raced, males were more likely to place poorly the longer it had been since they last raced. There was no significant difference between males and females in latency since the previous race. The effect of this interaction on performance is intriguing, but small and difficult to interpret. The significant negative correlation between mean eye temperatures before the races and latency since the previous race is at odds with a more intense stress response to the race-meet environment after longer rest periods as a potential explanation. It is possible that increasing latency since the previous race compromises race fitness, as extant data on racehorses shows an increase in the likelihood of sustaining a serious injury during a race with increasing days since last racing [35]. It is also possible that anticipation of racing is diminished by the lack of recent associations with track-related stimuli if the dog has not raced for more than a week or so, and this compromises performance. This is consistent with the current finding of a negative correlation with mean eye temperatures before the races.

Why latency since the previous race should affect male performance more than female performance is unclear and, to the authors’ knowledge, has not previously been reported in animal performance studies. As always, it is possible this reflects a statistical anomaly, and that simply increasing the sample size would verify the strength of this relationship. There was no significant difference in mean eye temperatures between sexes before or after the race and, to the authors’ knowledge, no sex differences in response to arousal have previously been reported in dogs. That said, there may be differences in how individuals of either sex behave in response to different levels of arousal.

### Catching pens

One of the goals of the current study was to investigate whether greyhounds in NSW races were being sufficiently rewarded for racing despite being unable to access the lure at the conclusion of races. We may assume that if greyhounds are finishing races without any penalty for failure to chase the lure, they are being sufficiently rewarded for racing at the time of observing them race. However, recording one race per greyhound cannot demonstrate that greyhounds are being sufficiently rewarded to continue racing indefinitely. As such, we searched for signs of frustration in the catching pen upon conclusion of the race. Frustration has been associated with increased aggression in dogs [36–38], which in turn, may manifest as an increased risk of attracting a penalty for marring during the race. It is also likely that what occurs in the catching pen influences a greyhound’s emotional associations with racing in general, and their willingness to enter the catching pen. For the purposes of the current study, the putative behavioural signs of frustration in the catching pen included jostling another dog, focusing on the lure gate, and changing directions (included only if the dog initiated the direction change rather than following another dog that had changed direction). We found these behaviours in 59.1% of greyhounds. The prevalence of these behaviours is concerning for two reasons. Firstly, it suggests that, at the end of the race, many greyhounds are still focused on chasing the lure when that opportunity is taken from them. If they are going to change direction, greyhounds almost always do so along the inside fence of the track or catching pen closest to the path of the lure arm, and often do so by orienting the head towards the lure’s direction of travel. The lure in transit makes a loud noise and, after passing the catching pen, may traverse more than half the track before coming to a halt, so the greyhounds can still hear it while, in the catching pen, unable to chase it. Orienting towards the lure gate indicates their focus on where they last saw the lure.

### Frustration in catching pens

The jostling that occurred after nearly every race in the current study is not necessarily an aggressive expression of frustration. However, the redirection of aggression towards conspecifics is a recognised consequence of being thwarted in obtaining a goal [see 39]. Other possible causes for this behaviour may be that it is playful in nature, or it may be a product of up to 8 galloping dogs coming to a halt together in a relatively small space. The dogs behind those in the lead may take a moment to react to the deceleration of dogs in front of them, resulting in bunching. Nonetheless, the indications that the greyhounds are still fixated on the lure raise the prospect that they are not disengaging from their goal even though they are unable to continue pursuing it. If their goal is to capture the lure, it may be less important whether they are successful or not and more important that they are not consistently thwarted in the process, but are instead assisted in disengaging from pursuit. The effects of frustration on behaviour may include heightened arousal as well as depressive disappointment [39]. Even if only a small number of greyhounds encounter the most detrimental effects of frustration in the catching pen, they may be subject to reduced interest in pursuing the rewards in question; an established response to frustrated non-reward [40].

### Teasers in catching pen

Only 11.3% of greyhounds at Richmond track were observed showed direct interest in the teasers in the catching pen. In contrast, 59.1% of greyhounds across all racetracks showed unresolved behaviours in the catching pen, and this was more than 3 times higher at the Wentworth Park and Gosford racetracks than at the Richmond racetrack. The distinctly lower incidence of unresolved behaviours in the catching pen at Richmond racetrack may be a result of the teasers in the catching pen, even though only a small portion of the racing greyhound population racing at Richmond showed active interest in the teasers. Taken together with the track effects, it appears Richmond racetrack is, in general, a less arousing environment than the other two racetracks. It is conceivable that this can be largely attributed to the presence of teasers, and this prospect warrants further investigation.

### Track effects

There was a significant difference among mean eye temperatures, both before and after races, at different tracks. This between-track difference was strongest with eye temperature before races, which is the metric most directly reflective of stressors arsing in the general racetrack environment. A general linear model, with mean eye temperatures before races e as the independent variable, revealed a significant positive influence of both Wentworth Park and Gosford compared to Richmond, with Richmond being associated with the lowest mean eye temperatures before the race. This may have been influenced by ambient temperature, but the effect of ambient temperature was neither strong nor significant. This suggests that some attributes of the Gosford racetrack may be inherently more stressful than Wentworth Park, and that those of Richmond may be less stressful than those of Wentworth Park. It is unclear why this may be. It may relate to the design of the kennel facilities, or perhaps even to the operational behaviour of personnel at the track and how handlers manage and interact with the dogs, or how quickly they enter the catching pen and restrain the dogs, or it may be the tracks are particularly attractive to owners of greyhounds that are more or less prone to distress. As noted in the previous section, this effect may also to some extent be attributable to the presence of teasers in the catching pen. Further investigation into which track attributes may influence greyhound stress before racing is important for the integrity of the sport so that vetting, kennelling, and pre-stir-up procedures can be designed to support greyhounds equitably.

There were significantly more behaviours indicative of arousal in the stir-up at Wentworth Park compared to Gosford. This may reflect the effects of a track design factor or a operational factor. The track at Wentworth Park has its catching pen adjacent to its stir-up pen and flush against the track itself, and directly in front of the kennels. In contrast, at Gosford, the stir-up pen is a considerable distance from the catch-pen, and even farther from the kennels. Richmond has the greatest distance between the stir-up pen and the track by a small margin, with the catching pen separated, and the kennels behind the stir-up pen. Handlers may also behave differently between tracks and may encourage more behaviours indicative of arousal at some tracks more than at others. This may reflect the handlers awareness of the available prize money or prestige of the races, the importance of a particular day of the week on which the races are held, or it may be entirely sub-conscious.

## Conclusions

The need to understand behavioural wastage in racing greyhounds is clear. Ambient temperature, race number, age and experience, start box, sex of dog, (frustration and teasers) in catching pens and track effects all affect the performance of greyhounds on tracks.

This is the first published study t of racing greyhound behaviour at race-meets and the first, for any racing code, to use IRT at racetracks to assess arousal. The results describe modest relationships between eye temperature and performance, but these effects are significant and may assist in the development of more detailed studies to identify specific factors that compromise performance and establish how they can be modified to reduce their negative effects. IRT before races may be more revealing than IRT after races due to the influence of core body temperature that may reflect the legacy effects of physical effort more than arousal, and the influence of ambient temperature. Attempts to use IRT before races bring significant timing and logistical challenges, but this study showed there is promise in eye temperature measurements before races to reveal the effects of experience at racetracks in the long- and short-term on behaviour and performance. For clearer results, it would be worthwhile investigating how IRT images may be collected closer to the races

This study also offers insights into how individuals within the racing greyhound population respond differently to the anticipation of racing, and how this might correlate with their preferences for rewards (e.g. teaser or lure) upon conclusion of any given race. Clearly, the use of teasers in the catching pen and the track effects on behaviour and arousal are both areas that merit further research to makes race-meets optimally arousing for racing greyhounds, and to improving reward availability and disengagement from the lure at the end of racse. The large percentage of dogs t showing signs of frustration and continuing to search for the lure when in the catching pen raises concern for both the physical and emotional wellbeing of greyhounds. The catching pen system has been in operation for many years, but may not support all racing greyhounds equitably in avoiding wastage. The period during which greyhounds are kennelled before their race may contribute to their stress at race-meets, so any means by which kennelling time can be reduced, especially for greyhounds in races late in the days’ program, are worth further investigation and suitable trials.

---

**S1 Figure Text:** Sketches of the three racing tracks in the study with kennels (a), stir-up yard (b) and catching pen (c) marked to show relative location of track features.

## References

1. McHugh M. Special Commission of Inquiry into the Greyhound Racing Industry in New South Wales. State of NSW; 2016 Jun pp. 1–269. Report No.: 1.

2. Starling MJ, McGreevy PD. Surveys on racing greyhound training practices in Australia. Sydney, Australia: Greyhound Racing NSW; 2017 Mar.

3. Greyhounds Australasia. Greyhounds Australasia Rules. 2016. pp. 1–75.

4. Cobb ML, Branson N, McGreevy PD, Bennett PC, Rooney NJ, Magdalinski T, et al. REVIEW & ASSESSMENT OF BEST PRACTICE. Working Dog Alliance Australia; 2015 Aug pp. 1–163.

5. Hutson GD, Haskell MJ. Pre-race behaviour of horses as a predictor of race finishing order. APPL ANIM BEHAV SCI. 1997;53: 231–248. doi:10.1016/S0168-1591(96)01162-8

6. Noteboom JT, Fleshner M, Enoka RM. Activation of the arousal response can impair performance on a simple motor task. J Appl Physiol. 2001;91: 821–831. doi:10.1152/jappl.2001.91.2.821

7. Roets A, Van Hiel A. An Integrative Process Approach on Judgment and Decision Making: The Impact of Arousal, Affect, Motivation, and Cognitive Ability. The Psychological Record. 2011;61: 11.

8. Travain T, Colombo ES, Heinzl E, Bellucci D, Previde EP, Valsecchi P. Hot dogs: Thermography in the assessment of stress in dogs (Canis familiaris)-A pilot study. J VET BEHAV. Elsevier Inc; 2015;10: 17–23. doi:10.1016/j.jveb.2014.11.003

9. Bouwknecht AJ, Olivier B, Paylor RE. The stress-induced hyperthermia paradigm as a physiological animal model for anxiety: A review of pharmacological and genetic studies in the mouse. NEUROSCI BIOBEHAV REV. 2007;31: 41–59. doi:10.1016/j.neubiorev.2006.02.002

10. de Lima V, Piles M, Rafel O, López-Béjar M, Ramón J, Velarde A, et al. Use of infrared thermography to assess the influence of high environmental temperature on rabbits. Research in Veterinary Science. Elsevier Ltd; 2013;95: 802–810. doi:10.1016/j.rvsc.2013.04.012

11. Dai F, Cogi NH, Heinzl EUL, Costa ED, Canali E, Minero M. Validation of a fear test in sport horses using infrared thermography. J VET BEHAV. Elsevier Inc; 2015;10: 128–136. doi:10.1016/j.jveb.2014.12.001

12. Bartolomé E, Sánchez MJ, Molina A, Schaefer AL, Cervantes I, Valera M. Using eye temperature and heart rate for stress assessment in young horses competing in jumping competitions and its possible influence on sport performance. Animal. Cambridge University Press; 2013;7: 2044–2053. doi:10.1017/S1751731113001626

13. Valera M, Bartolomé E, Sánchez MJ, Molina A, Cook N, Schaefer A. Changes in Eye Temperature and Stress Assessment in Horses During Show Jumping Competitions. Journal of Equine Veterinary Science. Elsevier Inc; 2012;32: 827– 830. doi:10.1016/j.jevs.2012.03.005

14. Stewart M, Stratton RB, Beausoleil NJ, Stafford KJ, Worth GM, Waran NK. Assessment of positive emotions in horses: Implications for welfare and performance. J VET BEHAV. Elsevier Inc; 2011;6: 296. doi:10.1016/j.jveb.2011.05.014

15. Fenner K, Yoon S, White P, Starling M, McGreevy P. The Effect of Noseband Tightening on Horses’ Behavior, Eye Temperature, and Cardiac Responses. Munderloh UG, editor. PLOS ONE. 2016;11: e0154179–20. doi:10.1371/journal.pone.0154179

16. Travain T, Colombo ES, Grandi LC, Heinzl E, Pelosi A, Previde EP, et al. How good is this food? A study on dogs’ emotional responses to a potentially pleasant event using infrared thermography. PHYSIOL BEHAV. Elsevier Inc; 2016;159: 80–87. doi:10.1016/j.physbeh.2016.03.019

17. Zanghi BM. Eye and Ear Temperature Using Infrared Thermography Are Related to Rectal Temperature in Dogs at Rest or With Exercise. FRONT VET SCI. 2016;3: R180–9. doi:10.3389/fvets.2016.00111

18. Riemer S, Assis L, Pike TW, Mills DS. Dynamic changes in ear temperature in relation to separation distress in dogs. PHYSIOL BEHAV. Elsevier Inc; 2016;167: 86–91. doi:10.1016/j.physbeh.2016.09.002

19. Part CE, Kiddie JL, Hayes WA, Mills DS, Neville RF, Morton DB, et al. Physiological, physical and behavioural changes in dogs (Canis familiaris) when kennelled: Testing the validity of stress parameters. PHYSIOL BEHAV. Elsevier Inc; 2014;133: 260–271. doi:10.1016/j.physbeh.2014.05.018

20. Gillette RL, Angle TC, Sanders JS, DeGraves FJ. An evaluation of the physiological affects of anticipation, activity arousal and recovery in sprinting Greyhounds. APPL ANIM BEHAV SCI. Elsevier B.V; 2011;130: 101–106. doi:10.1016/j.applanim.2010.12.010

21. Broom D. The welfare of livestock during road transport. In: Appleby MC, Cussen V, Garcés L, Lambert LA, Turner J, editors. Long Distance Transport and the Welfare of Farm Animals. London, UK: books.google.com; 2008. pp. 157–181.

22. Leadon D, Mullins E. Relationship between kennel size and stress in greyhounds transported short distances by air. VET REC. 1991;129: 70–73. doi:10.1136/vr.129.4.70

23. Bergeron R, Scott SL, Émond JP, Mercier F, Cook NJ, Schaefer AL. Physiology and behavior of dogs during air transport. The Canadian Journal of Veterinary Research. 2002;: 211–216.

24. CustomWeather. Syndicated Content for complete global weather coverage [Internet]. 2019 [cited 27 Feb 2019]. Available: https://customweather.com

25. Edgar JL, Nicol CJ, Pugh CA, Paul ES. Surface temperature changes in response to handling in domestic chickens. PHYSIOL BEHAV. Elsevier Inc; 2013;119: 195–200. doi:10.1016/j.physbeh.2013.06.020

26. Stewart M, Stafford KJ, Dowling SK, Schaefer AL, Webster JR. Eye temperature and heart rate variability of calves disbudded with or without local anaesthetic. PHYSIOL BEHAV. 2008;93: 789–797. doi:10.1016/j.physbeh.2007.11.044

27. Rizzo M, Arfuso F, Alberghina D, Giudice E, Gianesella M, Piccione G. Monitoring changes in body surface temperature associated with treadmill exercise in dogs by use of infrared methodology. Journal of Thermal Biology. Elsevier Ltd; 2017;69: 64–68. doi:10.1016/j.jtherbio.2017.06.007

28. Anthenill LA, Stover SM, Gardner IA, Hill AE. Risk factors for proximal sesamoid bone fractures associated with exercise history and horseshoe characteristics in Thoroughbred racehorses. AJVR. 2007;68: 760–771. doi:10.2460/ajvr.68.7.760

29. Leleu C, Cotrel C, Courouce-Malblanc A. Relationships between physiological variables and race performance in French standardbred trotters. VET REC. 2005;156: 339–342.

30. Gramm M, Marksteiner R. The Effect of Age on Thoroughbred Racing Performance. J Equine Sci. 2010;21: 73–79.

31. Takahashi T. The effect of age on the racing speed of Thoroughbred racehorses. J Equine Sci. Japanese Society of Equine Science; 2015;26: 43–48. doi:10.1294/jes.26.43

32. NSW Greyhound Breeders, Owners and Trainers’ Association LTD. Wentworth Park Greyhounds. In: Wentworth Park Greyhounds [Internet]. [cited 9 Feb 2018]. Available: http://www.wentworthpark.com.au/racing/how-to-bet-on-greyhounds

33. Dobbin A. Winning Box Stats… By The Numbers. In: The Greyhound Recorder [Internet]. 20 Dec 2018 [cited 28 Aug 2019]. Available: https://www.thegreyhoundrecorder.com.au/news/winning-box-stats-by-the-numbers-21219

34. Eager D, Hayati H, Mahdavi F, Hossain MI, Stephenson R, Thomas N. Identifying optimal greyhound track design for greyhound safety and welfare-Phase II-Progress Report-1 January 2016 to 31 December 2017. 2018.

35. Hernandez CE, Hinch G, Lea J, Ferguson D, Lee C. Acute stress enhances sensitivity to a highly attractive food reward without affecting judgement bias in laying hens. APPL ANIM BEHAV SCI. Elsevier B.V; 2015;163: 135–143. doi:10.1016/j.applanim.2014.12.002

36. Borchelt PL. Aggressive behavior of dogs kept as companion animals: Classification and influence of sex, reproductive status and breed. Applied Animal Ethology. Elsevier; 1983;10: 45–61. doi:10.1016/0304-3762(83)90111-6

37. Reisner IR. Differential diagnosis and management of human-directed aggression in dogs. VET CLIN NORTH AM SMALL ANIM PRACT. 2003;33: 303–320.

38. Luescher AU, Reisner IR. Canine Aggression Toward Familiar People: A New Look at an Old Problem. Veterinary Clinics of North America: Small Animal Practice. 2008;38: 1107–1130. doi:10.1016/j.cvsm.2008.04.008

39. Klinger E, 1975. Consequences of commitment to and disengagement from incentives. Psychological Review. 1975;82.

40. Carver CS. Negative Affects Deriving From the Behavioral Approach System. Emotion. 2004;4: 3–22. doi:10.1037/1528-3542.4.1.3

